# Calcium-dependent protein acyltransferase DHHC21 controls activation of CD4^+^ T cells

**DOI:** 10.1101/2020.09.01.277947

**Authors:** Bieerkehazhi Shayahati, Ying Fan, Savannah West, Ritika Tewari, Junsuk Ko, Tingting Mills, Darren Boehning, Askar M. Akimzhanov

## Abstract

Despite the recognized significance of reversible protein lipidation (S-acylation) for T cell receptor signal transduction, the enzymatic control of this post-translational modification in T cells remains poorly understood. Here, we demonstrate that DHHC21, a member of the DHHC family of mammalian protein acyltransferases, mediates agonist-induced S-acylation of proximal T cell signaling proteins. Using Zdhhc21^dep^ mice expressing a functionally deficient version of DHHC21, we show that DHHC21 is a calcium/calmodulin-dependent enzyme critical for activation of naïve CD4^+^ T cells in response to T cell receptor stimulation. We found that disruption of the calcium/calmodulin binding domain of DHHC21 does not affect thymic T cell development but prevents differentiation of peripheral CD4^+^ T cells into Th1, Th2, and Th17 effector T helper lineages. Our findings identify DHHC21 as an essential component of the T cell receptor signaling machinery and define a new role for protein acyltransferases in regulation of T cell-mediated immunity.

**Significance:** This study identifies DHHC21, a member of the DHHC family of mammalian protein acyltransferases, as a novel component of the TCR signaling pathway and demonstrates that this enzyme critically regulates activation and differentiation of CD4^+^ T cells by mediating rapid TCR-induced S-acylation of signaling proteins. This finding shows that protein acyltransferases can play a vital function in regulation of T cell-mediated immunity and thus serve as potential drug targets in diseases associated with altered immune system homeostasis.

## Introduction

CD4^+^ T cells are the master regulators of the adaptive immune response. During pathogen invasion, naïve CD4^+^ T cells become activated and differentiate into effector T helper (Th) lineages (1). Activation of CD4^+^ T cells is initiated upon engagement of the T cell receptor (TCR) by the antigen/major histocompatibility complex on the surface of the antigen presenting cell (2, 3). Ligation of the TCR by a foreign antigen triggers activation of Src-family kinases Lck and Fyn leading to phosphorylation of the TCR-associated CD3ζ-chain (2, 4). Phosphorylated TCR-CD3 complex amplifies the initial antigenic signal through recruitment of another regulatory kinase, ZAP-70 (5). Activated ZAP-70 proceeds to phosphorylate its downstream targets, primarily membrane-bound scaffolding proteins LAT and SLP-76 (6). Phosphorylated LAT and SLP-76 nucleate assembly of the multiprotein signaling complex termed “signalosome”. Formation of the signalosome further propagates the TCR signaling pathway resulting in activation of the important downstream mediators, such as phospholipase C-γ1 (PLC-γ1) and, cytoplasmic calcium influx, and, ultimately, leads to the transcriptional responses associated with activation and clonal expansion of CD4^+^ T cells (5, 7–9).

A number of proteins critically involved in regulation of the TCR signaling pathway were found to be S-acylated. Protein S-acylation (also known as S-palmitoylation) is a post-translational modification of cysteine thiols with long-chain fatty acids (10). Kinases Lck and Fyn were among the first mammalian proteins identified as S-acylated and this modification has been demonstrated to be necessary for their signaling function in T cells (11–14). Transmembrane adaptor LAT has been reported to be dually S-acylated at cysteine residues located proximally to the inner face of the plasma membrane, and a selective defect in LAT S-acylation has been found to be associated with a state of T cell functional unresponsiveness known as T cell anergy (15). Recently, a candidate-based screening of the proximal TCR signaling components allowed us to detect previously uncharacterized S-acylation of adaptor protein GRB2, PLC-γ1 and tyrosine kinase ZAP-70 (16, 17). A functional analysis of ZAP-70 S-acylation revealed that this modification is essential for ZAP-70 interaction with its protein substrates and subsequent transcriptional responses (17). Thus, S-acylation of T cell signaling proteins is emerging as an important regulatory mechanism controlling T cell activation and function. However, the enzymatic control of protein S-acylation in T cells remains enigmatic.

S-acylation of the mammalian proteins is catalyzed by a family of 23 protein acyltransferases with a common DHHC (Asp-His-His-Cys) motif within the catalytic core and, typically, four transmembrane domains (18). In our previous studies, we identified a member of this family, DHHC21, as a protein acyltransferase mediating S-acylation of several proximal T cell signaling proteins (16). Downregulation of DHHC21 in EL4 mouse lymphoma cells resulted in their unresponsiveness to TCR stimulation, suggesting that DHHC21 can be a critical regulator of the T cell effector function (16). To evaluate the contribution of DHHC21 to T cell-mediated immunity in an animal model, we used Zdhhc21^dep^ mice expressing a functionally deficient DHHC21 carrying an in-frame mutation resulting in a loss of a phenylalanine in the position 233 (ΔF233) (19, 20). We found that deletion of F233 disrupts TCR-induced calmodulin binding to DHHC21 indicating that DHHC21 is a calcium-regulated enzyme. Using acyl-resin assisted capture (Acyl-RAC) technique, we found that the ΔF233 mutant of DHHC21 was unable to mediate TCR-induced S-acylation of critical T cell signaling proteins resulting in diminished activation of the TCR pathway. Although expression of the ΔF233 mutant did not affect T cell development, naïve CD4^+^ cells isolated from peripheral lymphoid organs of Zdhhc21^dep^ mice exhibited decreased activation of the proximal TCR signaling components and suppressed cytoplasmic calcium release in response to TCR stimulation. Consequently, Zdhhc21^dep^ CD4^+^ T cells showed significantly decreased upregulation of the surface activation marker CD69 and reduced production of the pro-inflammatory cytokine interleukin-2 (IL-2) upon extended stimulation of the TCR. Furthermore, the impaired DHHC21 function prevented naïve Zdhhc21^dep^ CD4^+^ T cells from differentiation into effector T helper subtypes, suggesting that protein acyltransferase DHHC21 plays an important role in peripheral T cell immunity.

## Results

### F233 is required for calcium-dependent calmodulin binding to DHHC21

The spontaneous depilated (“dep”) mutation of DHHC21 (Zdhhc21^dep^, MGI:94884) was initially described as a recessive mutation on chromosome 4 characterized by variable hair loss and abnormal hair structure and was later mapped to the *zdhhc21* gene (19, 20). Delayed follicle differentiation, endothelial dysfunction and reduced vascular tone observed in Zdhhc21^dep^ mice (19, 21, 22) suggested that deletion of F233 causes a functional deficiency of DHHC21. However, the molecular mechanism underlying dysfunction of the ΔF233 DHHC21 mutant remained unresolved.

Our analysis of the primary DHHC21 sequence revealed that F233 is located within a putative 1-14 calcium/calmodulin-binding motif at the C-terminal cytoplasmic tail of the enzyme (**Fig. 1A**). To experimentally validate calmodulin binding to DHHC21, we performed the co-immunoprecipitation assay using CD4^+^ T cells isolated from WT and Zdhhc21^dep^ mice. To determine whether DHHC21 interacts with calmodulin in a calcium-dependent manner, we treated cells with thapsigargin (TG), a potent SERCA pump inhibitor (23), to invoke a cytoplasmic calcium influx form the ER stores. Endogenously expressed DHHC21 was then immunoprecipitated from the cell lysates and analyzed by immunoblotting. As shown in **Fig 1B**, treatment of WT CD4^+^ T cells with 5 µM TG resulted in co-immunoprecipitation of DHHC21 with calmodulin indicating the presence of a functional calcium/calmodulin binding domain. However, calmodulin was not detected in DHHC21 immunoprecipitates from Zdhhc21^dep^ CD4^+^ T cells suggesting that deletion of F233 disrupts binding between DHHC21 and calmodulin (**Fig 1B**). Since calmodulin is known to relay calcium signals through direct binding to its targets, this result suggests that the ΔF233 mutation could prevent activation of the enzyme in response to TCR-induced calcium release, thus leading to the functional deficiency of DHHC21.

**Fig. 1.**
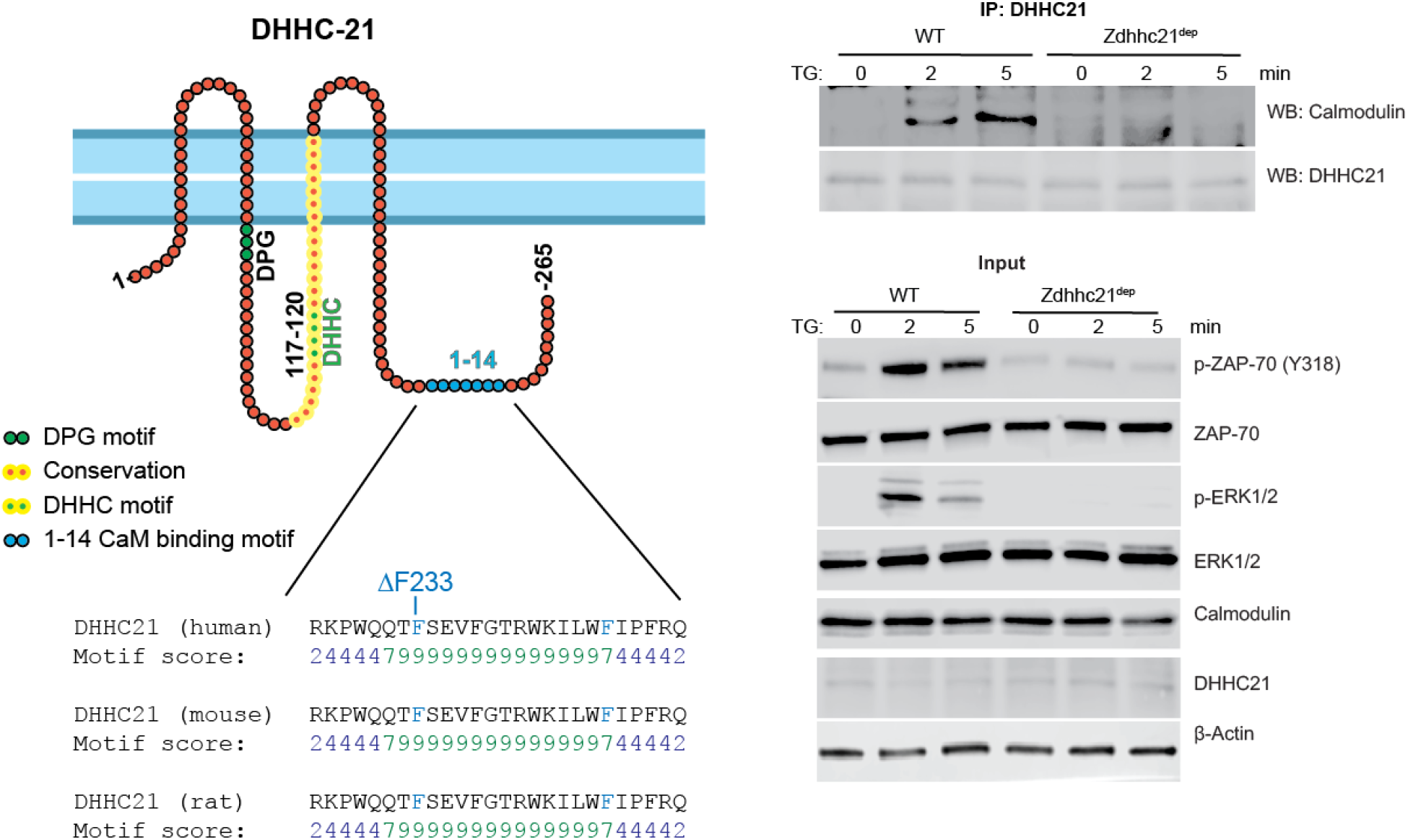
DHHC21 has a calcium/calmodulin binding motif. **(A)** Schematic depiction of DHHC21 illustrating the approximate localization of the predicted 1-14 calcium/calmodulin binding motif within the C-terminal cytoplasmic tail. The 1-14 motif is completely conserved between species. Location of the “dep” mutation (ΔF233) is indicated. **(B)** Calcium-dependent co-immunoprecipitation of DHHC21 and calmodulin. CD4^+^ T cells from WT or Zdhhc^dep^ mice were treated with 10µM thapsigargin (TG) to evoke calcium release and DHHC21 was immunoprecipitated from the cell lysates. Binding of calmodulin was detected by immunoblotting.

### DHHC21 mediates TCR-induced S-acylation of T cell signaling proteins

We next sought to examine the effect of the ΔF233 mutation on acyltransferase activity of DHHC21. Previously, we detected direct stimulus-dependent interaction between DHHC21 and kinases Lck and Fyn (16). Furthermore, we showed that shRNA-mediated downregulation of DHHC21 significantly decreased S-acylation of Src-family kinases Lck and Fyn, and PLC-γ1 in resting EL4 lymphoma cells, suggesting that these proteins are direct targets of DHHC21 (16). To test whether ΔF233 DHHC21 is able to mediate S-acylation of its protein substrates, we isolated CD4^+^ T cells from spleen and lymph nodes of WT and Zdhhc21^dep^ mice and subjected them to the acyl-resin assisted capture (Acyl-RAC) assay (**Fig. 2A**) (24). Briefly, free thiol groups in cell lysates were blocked with S-methyl methanethiosulfonate (MMTS) and the thioester bond between cysteine residues and fatty acid moieties was selectively cleaved with neutral hydroxylamine (HA). The newly exposed cysteine thiol groups were then reacted with a thiol-reactive resin to capture the S-acylated proteins. As shown in **Fig. 2B**, deletion of F233 did not affect basal S-acylation levels of Lck, Fyn, PLC-γ1 and ERK1/2 proteins in quiescent T cells. In WT CD4^+^ T cells, short-term TCR stimulation with cross-linked anti-CD3/CD28 antibodies resulted in increased S-acylation of Lck, ERK1/2, and PLC-γ1. However, we did not observe TCR-induced S-acylation of these proteins in CD4^+^ T cells isolated from Zdhhc21^dep^ mice (**Fig. 2**). Our observations suggest that although ΔF233 DHHC21 can still support basal lipidation, a functional calcium/calmodulin binding domain is required to mediate antigen-induced S-acylation of DHHC21 protein substrates. These results were also consistent with previously reported substrate specificity of DHHC21 toward Lck, Fyn, and PLC-γ1 (16) and indicate that ERK1/2, recently identified as a novel S-acylated protein in T cells (data not shown), could be another target of DHHC21. No significant changes were observed in S-acylation of Fyn in stimulated T cells (**Fig. 2**), suggesting that a different protein acyltransferase could contribute to S-acylation of this kinase.

**Fig. 2.**
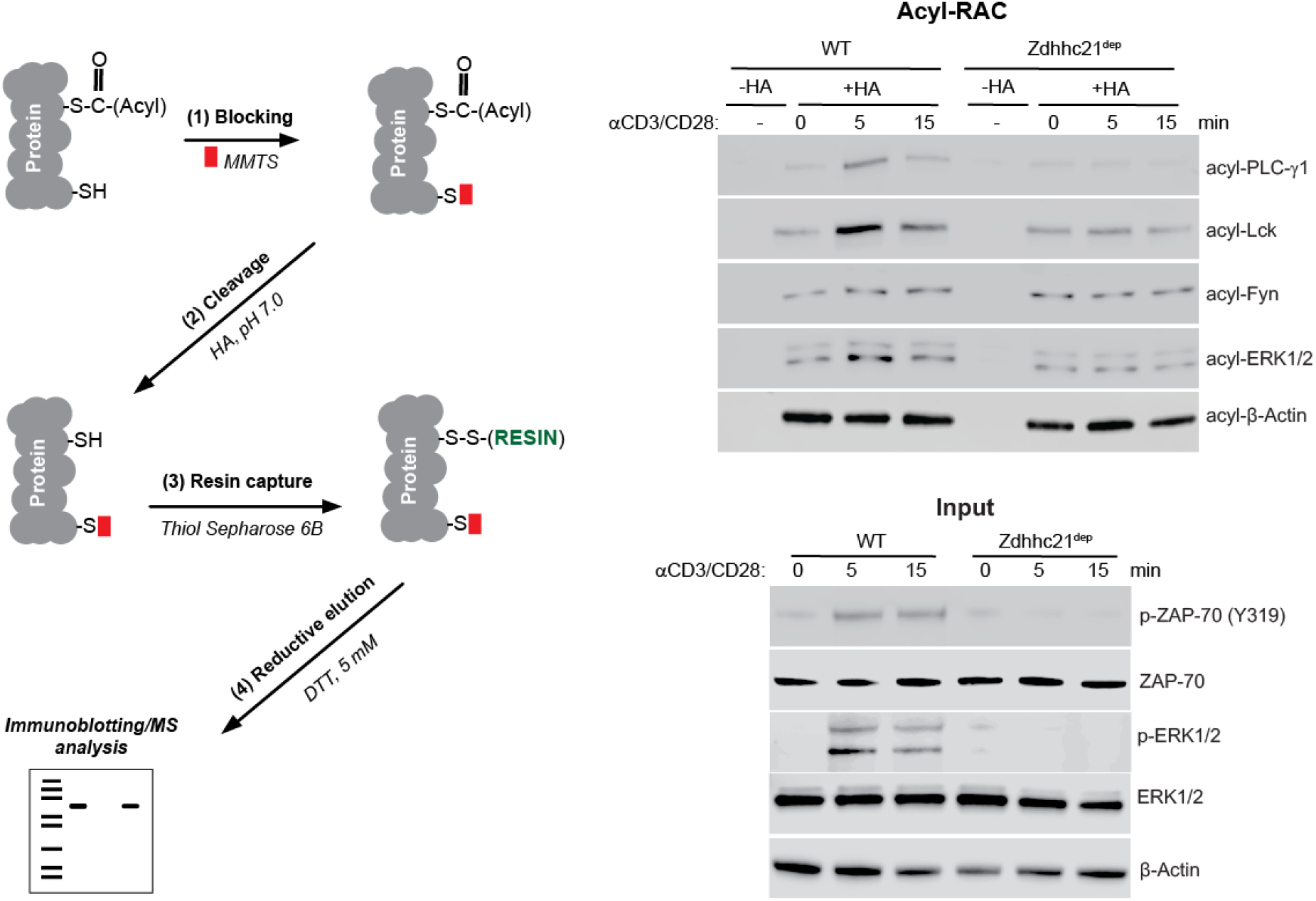
DHHC21 mediates TCR-induced S-acylation of signaling proteins. **(A)** Overview of the acyl-resin assisted capture (Acyl-RAC) assay. Free cysteine thiols (-SH) are irreversibly blocked by S-methyl methanethiosulfonate (MMTS) following cell lysis. Thioester bonds between cysteines and acyl groups are then specifically cleaved by neutral hydroxylamine (HA). The newly formed free thiol groups are captured by the thiol-reactive sepharose and S-acylated proteins detected by immunoblotting. **(B)** TCR-induced protein S-acylation. CD4^+^ T cells from WT or Zdhhc21^dep^ mice were stimulated with cross-linked anti-CD3/CD28 antibodies for indicated times and protein S-acylation (acyl-) was determined using Acyl-RAC assay. Samples not treated with hydroxylamine (-HA) were used as a negative control. β-Actin, a known S-acylated protein (32), was used as a loading control.

### DHHC21 is required for initiation of the TCR signaling pathway

Our data indicate that loss of F233 inhibited the ability of DHHC21 to S-acylate its targets in response to TCR stimulation. Since identified DHHC21 protein substrates are components of the proximal TCR pathway, we next aimed to determine whether the functional deficiency of ΔF233 DHHC21 affects initiation of the early TCR signaling events. CD4^+^ T cells purified from spleen and lymph nodes of WT and Zdhhc21^dep^ mice were stimulated with cross-linked anti-CD3/CD28 antibodies and phosphorylation of key TCR signaling proteins was analyzed by immunoblotting. We found that activation of the first signaling TCR kinase Lck, measured with phospho-Src family antibody, was diminished in Zdhhc21^dep^ CD4^+^ T cells (**Fig. 3A**). TCR-induced activation of critical downstream proteins ZAP-70, LAT, PLC-γ1, ERK1/2, JNK and p38 was also suppressed in these cells indicating that DHHC21 could be one of the earliest signal mediators in the TCR pathway.

**Fig. 3.**
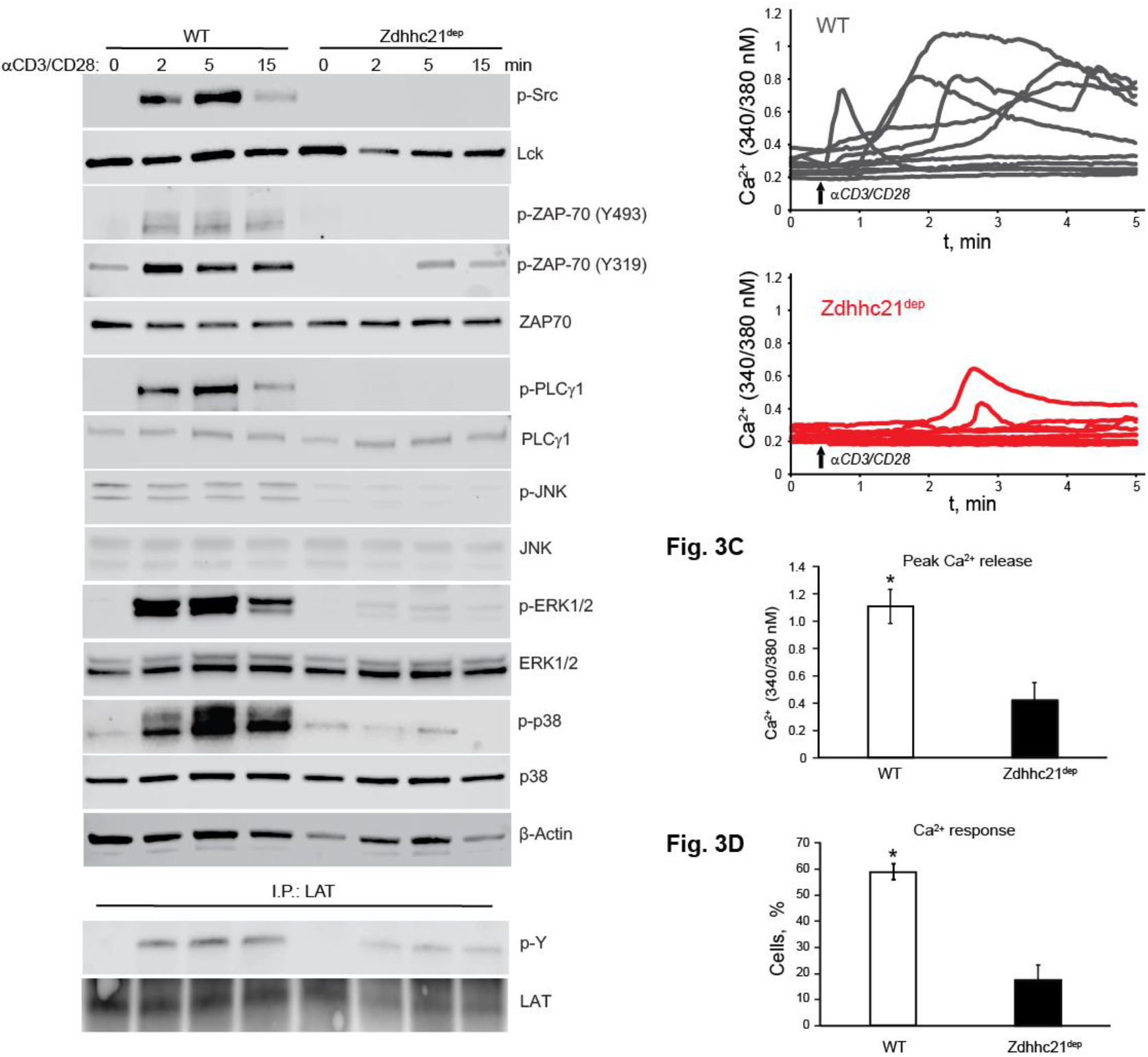
TCR signaling is impaired in Zdhhc21^dep^ CD4^+^ T cells. **(A)** CD4^+^T cells from WT or Zdhhc21^dep^ mice were stimulated with cross-linked anti-CD3/CD28 antibodies for indicated times and indicated phospho-proteins (p-) and total proteins were analyzed by immunoblotting. β-Actin was used as a loading control. The results shown are representative of three independent experiments. **(B)** CD4^+^ T cells were loaded with Fura-2 calcium indicator and pre-incubated with anti-CD3/CD28 antibodies. IgG antibody was added to induce cross-linking after an observation period (upward arrow). Shown are representative traces from single WT (gray) and Zdhhc21^dep^ (red) cells. **(C)** Peak calcium release. CD4^+^T cells were treated as described in (B). Data represent maximum calcium values averaged from four independent experiments (∼100 WT or Zdhhc21^dep^ cells per experiment). ± S.E.M. **(D)** Calcium response pattern. CD4^+^T cells were treated as described in (B). Cells showing at least 10% increase in calcium values upon TCR stimulation were identified as responders. Data represent averaged percentage of responders from four independent experiments (∼100 WT or Zdhhc21^dep^ CD4^+^T cells per experiment). ± S.E.M.

Calcium serves as a critical second messenger during T cell activation. The cytoplasmic calcium influx is triggered upon TCR-dependent activation of PLC-γ1 which generates inositol-1,4,5-trisphosphate required for rapid calcium release from the ER stores (25, 26). To further test the role of DHHC21 in initiation of the proximal TCR signaling cascade, we loaded WT and Zdhhc21^dep^ CD4^+^ T cells with Fura-2 calcium indicator and assessed changes in cytoplasmic calcium concentration upon T cell activation. Stimulation of the TCR with anti-CD3/CD28 antibodies evoked immediate calcium responses in ∼80% of the WT CD4^+^ T cells (**Fig. 3B, C**). Consistent with inhibited activation of PLC-γ1, we found that TCR engagement induced calcium release in only ∼20% of Zdhhc21^dep^ CD4^+^ T cells and the cytoplasmic calcium concentrations peaked at significantly reduced levels (**Fig. 3B-D**). This result indicates that fully functional DHHC21 is required to support normal calcium dynamics in stimulated T cells.

### Suppressed T cell activation in Zdhhc21^dep^ mice

Thus far, our results show that the ΔF233 mutation of DHHC21 causes disruption of the early TCR signaling events, including activation of regulatory kinases and calcium mobilization from the ER. To further investigate the role of DHHC21 in T cell activation, we stimulated CD4^+^ T cells isolated from Zdhhc21^dep^ mice and their WT littermates with plate-bound anti-CD3/CD28 antibodies for 24 hours and assessed induction of T cell activation markers. We found that in contrast to WT cells, activated Zdhhc21^dep^ CD4^+^ T cells showed markedly decreased surface expression of CD69 (**Fig. 4A, B**). Similarly, production of the pro-inflammatory cytokine IL-2 in response to TCR stimulation was significantly reduced in CD4^+^T cells expressing the ΔF233 DHHC21 mutant (**Fig. 4C**). Thus, consistent with previous observations, our data suggest that DHHC21 is required for T cell activation upon TCR engagement.

**Fig. 4.**
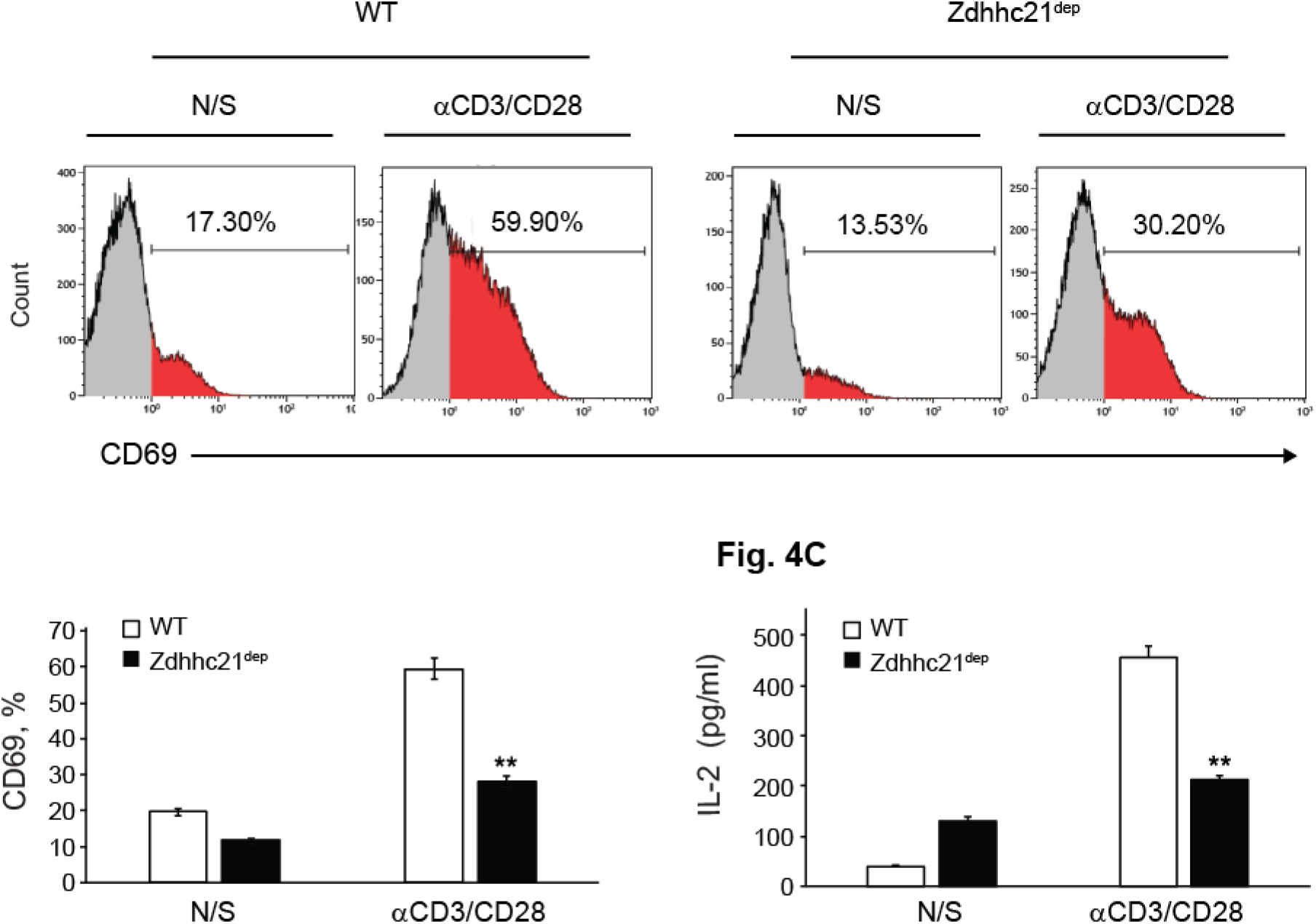
DHHC21 is required for T cell activation. **(A)** Surface expression of CD69 on CD4^+^ T cells from WT or Zdhhc21^dep^ mice. CD4^+^ T cells were stimulated with plate-bound anti-CD3/CD28 antibodies for 24 h and analyzed by flow cytometry. **(B)** Average percentage of CD69^+^ T cells from WT or Zdhhc21^dep^ mice. CD4^+^ T cells were stimulated with plate-bound anti-CD3/CD28 antibodies for 24 h and analyzed by flow cytometry. The graph shows mean ± SD values from three independent experiments. **, P < 0.01, Student t-test. **(C)** IL-2 production by CD4^+^ T cells from WT or Zdhhc21^dep^ mice. CD4^+^ T cells were stimulated with plate-bound anti-CD3/CD28 antibodies for 24 h and collected supernatants were assayed by ELISA. The graph shows mean ± SD values from three independent experiments. **, P < 0.01, Student t-test.

### F233 is dispensable for T cell development

To examine a possible role of DHHC21 in T cell development and differentiation, we assessed the frequency of major T cell subpopulations in 4-6 weeks old Zdhhc21^dep^ mice and their WT littermates in thymus and peripheral lymphoid organs. Flow cytometry analysis of double-negative (DN, CD4^−^ CD8^−^), double-positive (DP, CD4^+^CD8^+^) and single positive (SP, CD4^+^CD8^−^ and CD4^−^ CD8^+^) thymocytes revealed no difference in relative proportions of these populations (**Fig. 5A**). The total numbers of thymocytes and thymus size were also similar in WT and Zdhhc21^dep^ mice and no difference was detected in expression of CD44 and CD25 on the surface of the DN thymocytes (data not shown and **Fig. 5A, B**) suggesting that the ΔF233 DHHC21 mutation did not affect DN-to-DP and DP-to-SP transitions during T cell maturation.

**Fig. 5.**
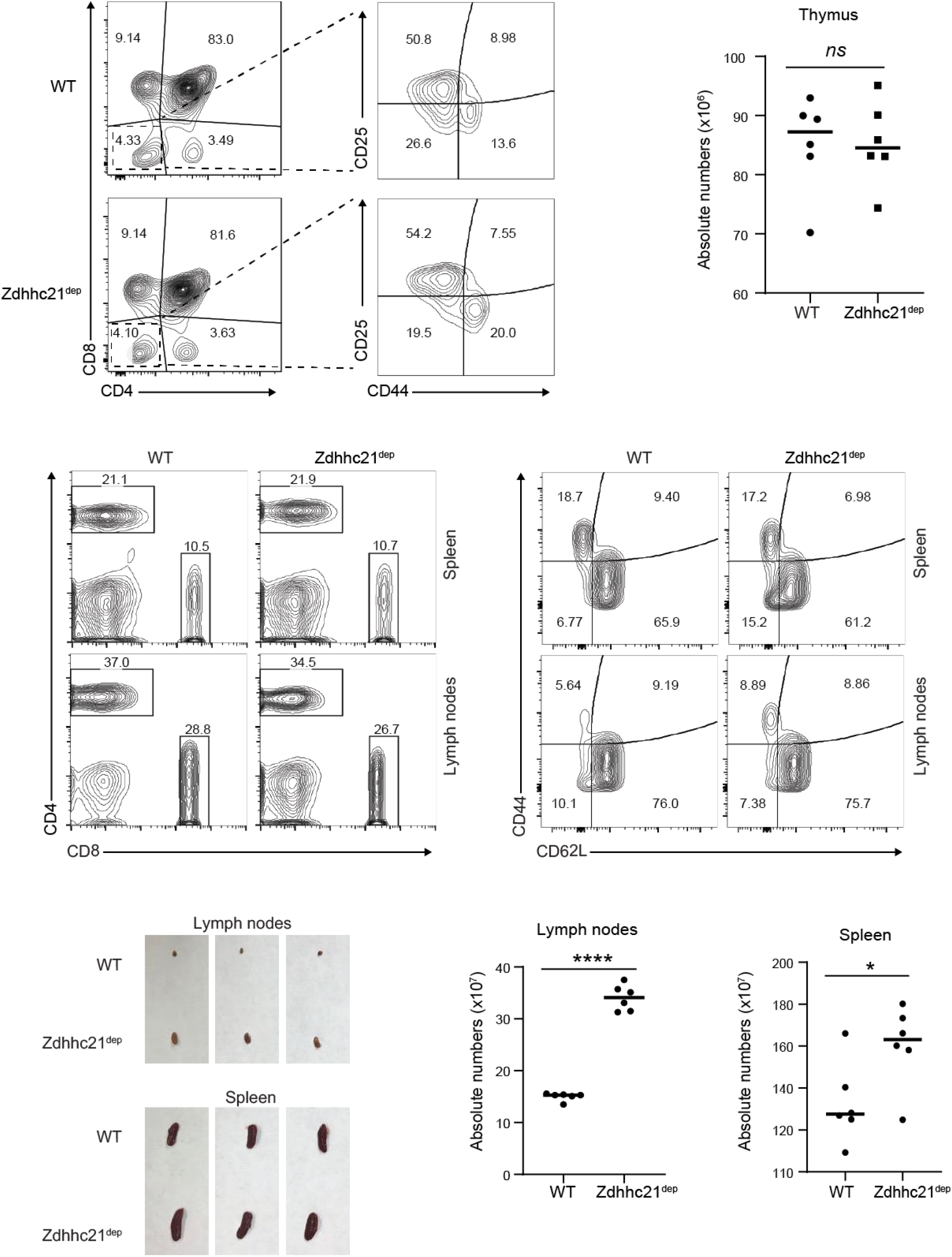
Normal thymic development in Zdhhc21^dep^ mice. **(A)** Flow cytometry analysis of cells isolated from thymus of WT and Zdhhc21^dep^ mice. Plots of CD4^+^ and CD8^+^ cells are gated on CD3^+^ cells. Plots of CD25^+^ and CD44^+^ cells are gated on CD4^-^CD8^-^ cells. **(B)** The absolute numbers of thymocytes from WT and Zdhhc21^dep^ mice. Cell numbers were calculated based on the relative percentages determined by flow cytometry analysis and total cell numbers from thymus. ns, P >0.05, Student t-test. **(C)** Flow cytometry analysis of cells isolated from spleen and lymph nodes of WT and Zdhhc21^dep^ mice. Plots of CD4^+^ and CD8^+^ cells are gated on CD3^+^ cells. **(D)** Flow cytometry analysis of naive (CD44^lo^CD62L^hi^) and effector memory (CD44^hi^CD62L^lo^) CD4^+^ T cells isolated from spleen and lymph nodes of WT and Zdhhc21^dep^ mice. **(E)** Peripheral lymphoid organs in WT and Zdhhc21^dep^ mice. **(F)** The absolute numbers of CD3^+^ cells isolated from spleen and lymph nodes of WT and Zdhhc21^dep^ mice. Cell numbers were calculated based on the relative percentages determined by flow cytometry analysis and total cell numbers from thymus. ****, P < 0.001 (left) and *, P < 0.05 (right), Student t-test.

The analysis of T cell populations in peripheral lymphoid organs also showed no substantial differences between CD4^+^ and CD8^+^ T cell percentages in WT and Zdhhc21^dep^ mice (**Fig. 5C**). Similarly, no changes were detected in populations of naïve (CD44^lo^CD62L^hi^) and effector memory (CD44^hi^CD62L^lo^) T cells (**Fig. 5D**). However, we noted the consistent increase in size of spleen and lymph nodes of Zdhhc21^dep^ mice accompanied by statistically significant increase in absolute cell numbers (**Fig. 5E, F**), indicating that the attenuated DHHC21 function could affect regulation of the peripheral T cell immunity.

### DHHC21 is important for peripheral CD4^+^ T cell immunity

Given the critical role of DHHC21 in TCR-mediated T cell activation, we hypothesized that expression of the functionally deficient ΔF233 DHHC21 mutant affects the ability of naïve CD4^+^ T cells to differentiate into functionally distinct effector T helper (Th) cell lineages. To test the hypothesis, we isolated naïve CD4^+^ T cells from spleen and lymph nodes of WT and Zdhhc21^dep^ mice and incubated them under Th1, Th2, or Th17 polarizing conditions. After 5 days, cells were re-stimulated with plate-bound anti-CD3/CD28 antibodies and production of lineage specific cytokines was assessed with ELISA. As shown in **Fig. 6A**, Zdhhc21^dep^ CD4^+^ T cells produced significantly less IFN-γ cytokine when cultured under Th1 conditions. Under Th2 polarizing conditions, Zdhhc21^dep^ CD4^+^ T cells failed to secrete IL-6 and IL-13 cytokines. Under Th17 conditions, we observed significantly decreased production of IL-17A in Zdhhc21^dep^ CD4^+^ T cells when compared to WT CD4^+^ T cells. Interestingly, IFN-γ production under Th17 conditions was increased in Zdhhc21^dep^ CD4^+^ T cells indicating a possible role of DHHC21 in Th17-mediated autoimmunity.

**Fig. 6.**
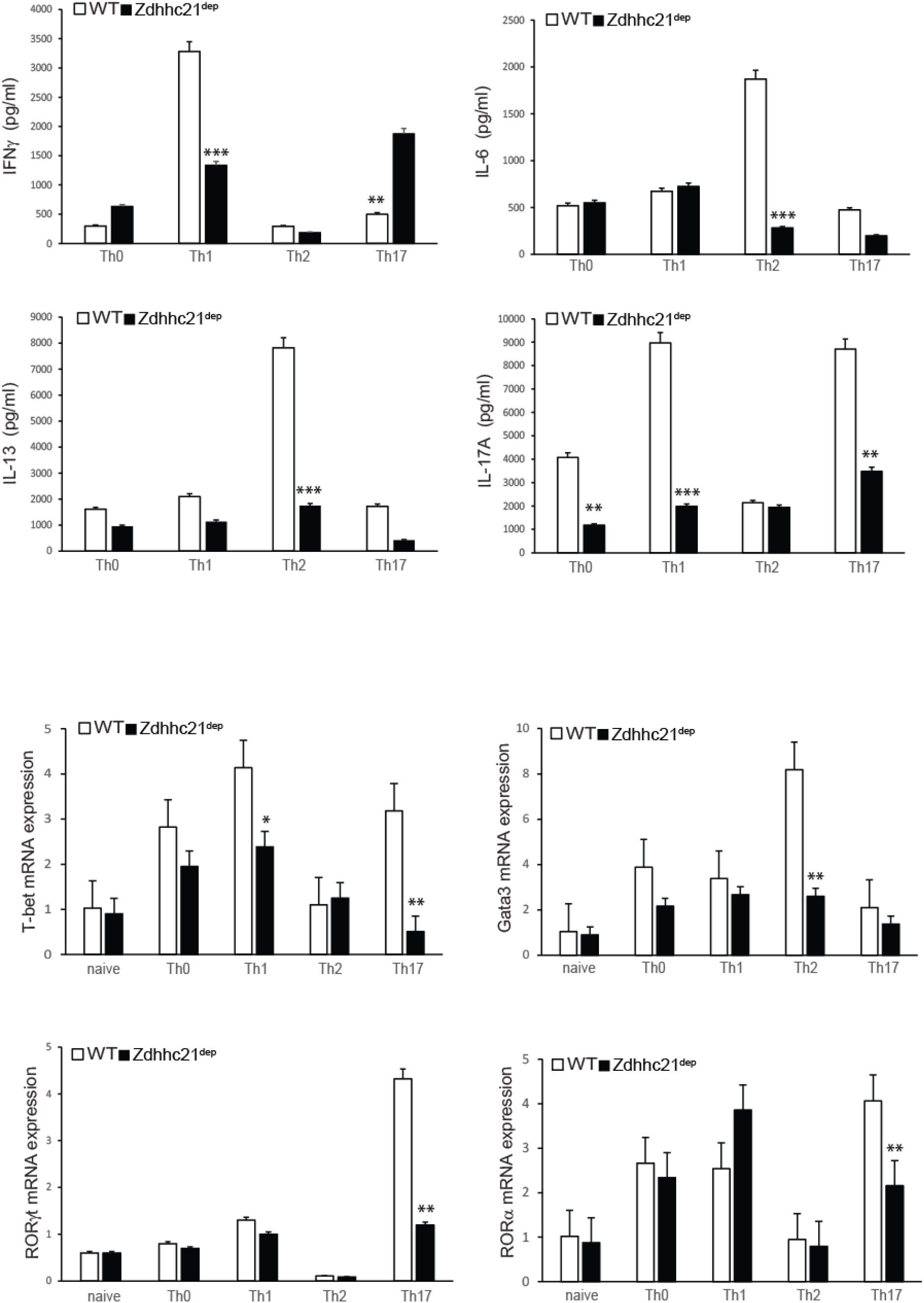
DHHC21 regulates CD4^+^ T activation and differentiation. Naïve CD4^+^ T cells from WT or Zdhhc21^dep^ mice were stimulated with plate-bound anti-CD3/CD28 antibodies and incubated under neutral (Th0) or polarizing (Th1, Th2, and Th17) conditions for 5 days, then CD4^+^ T cells were re-stimulated with plate-bound anti-CD3 antibodies for 6 h. **(A)** Effector cytokine production measured by ELISA. The graph shows mean ± SD values from at least three independent experiments. **(B)** mRNA expression of Th lineage-specific transcription factors determined by qRT-PCR normalized to 18S RNA. The graph shows mean ± SD values from at least three independent experiments. *, P < 0.05; **, P < 0.01; ***, P < 0.001, Student t-test.

To further examine the role of DHHC21 in effector T cell differentiation, we used real-time PCR to assess expression of lineage-specific transcription factors in CD4^+^ T cells polarized under Th1, Th2 or Th17 conditions. We found that in comparison to WT cells, CD4^+^ T cells from Zdhhc21^dep^ mice demonstrated significantly decreased expression of T-bet under Th1 and Th17 conditions, and failed to upregulate expression of GATA-3 under Th2 conditions (**Fig. 6B**). Likewise, upregulation of RORγτ and RORα, the signature Th17 genes, was significantly diminished in Zdhhc21^dep^ CD4^+^ T cells differentiated under Th17 conditions (**Fig. 6B**). Overall, our data indicate that DHHC21 is not only important for initial T cell activation but also required for proper differentiation of peripheral CD4^+^ T cells into major T helper lineages.

## Discussion

Several classes of regulatory enzymes cooperate to ensure proper activation of T cells in response to invading pathogens. Here, we report a previously uncharacterized role of protein acyltransferases – enzymes mediating reversible protein S-acylation – in regulation of the T cell function. Specifically, our study demonstrates that DHHC21, a member of the DHHC family of mammalian protein acyltransferases, is essential for TCR-dependent CD4^+^ T cell activation and subsequent differentiation into major T helper lineages.

The importance of protein S-acylation for the T cell immune responses became evident since the discovery that this post-translational modification critically affects the plasma membrane localization and function of well-characterized components of the canonical TCR signaling pathway, such as Lck, Fyn, LAT and, recently, ZAP-70 (11–15, 17, 27). These findings implied that the TCR signal transduction could be modulated by the enzymes catalyzing protein S-acylation, known as the DHHC family of protein acyltransferases (18). However, the exact identity of protein acyltransferases involved in TCR signaling, their regulatory mechanisms, and physiological significance remained largely unknown. In our previous study aimed at identification of the S-acylating enzyme for Lck, we found that protein acyltransferase DHHC21 expressed in EL4 lymphoma cell line has substrate preferences for Src-family kinases Lck and Fyn, and PLC-γ1, recently identified as S-acylated (16). Identification of key TCR signaling proteins as DHHC21 substrates suggested that this enzyme could be an important part of the proximal TCR signaling cascade. In this study, to further explore the role of the DHHC21 in regulation of TCR-induced T cell activation, we took advantage of the Zdhhc21^dep^ mouse strain in which a single phenylalanine residue (F233) within the C-terminal tail of DHHC21 is deleted (19).

We identified F233 as the first phenylalanine within a classical 1-14 calcium/calmodulin binding motif and found that deletion of this residue disrupts calcium-dependent interaction between calmodulin and DHHC21. Given a well-known role of calmodulin in mediation of intracellular calcium signals, this finding suggests that enzymatic activity of DHHC21 can be modulated by the cytoplasmic calcium influx triggered upon TCR engagement. Indeed, in our previous study we have determined that S-acylation of DHHC21 substrate Lck was dependent on PLC-γ1-mediated calcium release from the ER stores and that elimination of the calcium signal prevented agonist-induced changes in Lck S-acylation (28). Consistent with these observations, we found that disruption of the calcium/calmodulin binding motif of DHHC21 did not affect the basal S-acylation levels, but abrogated TCR-induced increase in S-acylation of DHHC21 protein targets. While more detailed studies are required to fully elucidate the mechanism of DHHC21 regulation, our findings indicate that similar to other components of the TCR pathway, increases in enzymatic activity of DHHC21 can be rapidly triggered by the TCR-induced cytoplasmic calcium influx.

Extremely rapid (within minutes) increase of protein S-acylation in response to agonist stimulation has been recently observed in several studies (16, 17, 28, 29). However, the physiological meaning of these changes remained unclear since conventional approaches to study the role of protein S-acylation rely on complete elimination of this modification through site-directed mutagenesis or pharmacological treatment. Our data suggests that the ΔF233 DHHC21 mutant preserved its basal enzymatic activity but was unable to respond to the TCR treatment by increasing S-acylation of its targets. Thus, the nature of the ΔF233 mutation provided us with a unique opportunity to examine the function of stimulus-dependent changes in S-acylation separately from the effects of S-acylation itself. We found that despite no detectable difference in basal protein S-acylation between WT and ΔF233 DHHC21 mice, CD4^+^ T cells isolated from Zdhhc21^dep^ strain failed to respond to TCR stimulation, as evidenced by diminished activation of the proximal TCR pathway components, suppressed calcium release and significantly reduced upregulation of major T cell activation markers. The fast kinetics of stimulus-induced protein S-acylation and its dramatic effect on T cell activation agree with a model in which DHHC21 is an integral part of the proximal TCR signaling machinery mediating the antigen signal transduction through rapid S-acylation of signaling proteins.

Initial T cell responses determine the long-term outcomes of TCR stimulation, such as development of immature T cells in the thymus and differentiation of naïve T cells in the peripheral lymphoid organs. We found that expression of ΔF233 DHHC21 had no apparent effect on populations of DN, DP and SP thymocytes suggesting that either DHHC21 is dispensable for T cell development or the reduced functionality of the ΔF233 mutant was sufficient to support DN-to-DP and DP-to-SP transitions during T cell maturation. However, we found that the inability of ΔF233 DHHC21 to respond to the TCR-induced calcium signals significantly impaired differentiation of naïve CD4^+^ T cells into Th1, Th2 and Th17 effector lineages. These findings suggest that DHHC21 plays a more prominent role in peripheral immune responses and warrants further studies to investigate a potential role of DHHC21 in adaptive immunity.

In summary, our study identifies protein acyltransferase DHHC21 as an essential component of the canonical TCR signaling pathway and demonstrates that this enzyme promotes early signal transduction by mediating rapid TCR-induced S-acylation of its protein targets. Our findings imply that protein acyltransferases, similar to other classes of regulatory enzymes can play a vital function in regulation of T cell-mediated immunity and thus serve as potential drug targets in diseases associated with altered immune system homeostasis.

## Material and Methods

### Antibodies and reagents

The following antibodies were purchased from Cell Signaling: Lck (Cat. 2787), p-Src (Cat. 2101), LAT (Cat.45533), Fyn (Cat.4023), p-Tyr-100 (Cat.9411), p-PLC-γ1 (Cat.8713), PLC-γ1 (Cat.2824), p-ZAP-70 (Cat. 2717), ZAP-70 (Cat. 3165), p-Erk1/2 (Cat. 4370), ERK1/2 (Cat. 4376), JNK (Cat.4668), p-JNK (Cat.9252). DHHC21 antibody was made in our laboratory. The following reagents were purchased from Sigma-Aldrich: Protein A Sepharose (Cat. p6649), Hydroxylamine (Cat. 55459), S-methyl methanethiosulfonate (MMTS) (Cat. 208795), n-Dodecyl β-D-maltoside (DDM) (Cat. D4641), Thiopropyl-Sepharose 6B (Cat. T8387), Poly-L-lysine (Cat. P8920), Phosphatase Inhibitor Cocktail 2 (Cat. P5726), Complete Protease Inhibitor Cocktail tablets (Cat. 11836170001). ML211 (Cat. 17630) was purchased from Cayman. Cell-stimulating mAbs were bought from BD Biosciences, San Diego, CA (anti-CD28), purified from a specific hybridoma (anti-CD3).

### Mice

Zdhhc21^dep^ mice were bred in our facility under specific pathogen-free conditions in accordance with the recommendations in the Guide for the Care and Use of Laboratory Animals of the National Institutes of Health. The animals were handled according to the animal care protocol (AWC 18-0131) approved by the Animal Welfare Committee (AWC), the Institutional Animal Care and Use Committee (IACUC) for the University of Texas Health Science Center at Houston (UTHealth).

### Western blot

For protein immunoblotting, after the indicated stimulation, cells were washed twice with cold PBS and then lysed in 1% DDM lysis buffer (1 % Dodecyl β-D-maltoside (DDM) in DPBS; 10 µM ML211; Phosphatase Inhibitor Cocktail 2 (1:100); Protease Inhibitor Cocktail (1X), PMSF (10 mM)).Protein concentration was determined by Bio-Rad Bradford protein assay and equal amounts of protein were loaded for immunoblot analysis. Lysates (30μg protein) were separated by the SDS-PAGE, and then transferred to NC membranes (BioRad). Membranes were blocked in 5% bovine serum albumin (BSA) in PBS-T (0.1% Tween-20 in PBS buffer) for 1 hr at room temperature, and then incubated with the indicated primary antibodies overnight at 4 °C. The membranes were then incubated with anti-mouse or anti-rabbit IgG conjugated with horseradish peroxidase at room temperature for 1 hr. After three washes in PBS-T, membranes were imaged using the LI-COR Odyssey Scanner (LI-COR Biosciences; Lincoln, NE). Brightness and contrast were adjusted in the linear range using the Image Studio software (LI-COR).

### Acyl-Resin Assisted Capture (Acyl-RAC) Assay

Protein S-acylation status was assessed by Acyl-RAC as described (24). Cells were lysed in 1% DDM lysis buffer (1 % Dodecyl β-D-maltoside (DDM) in DPBS; 10 µM ML211; Phosphatase Inhibitor Cocktail 2 (1:100); Protease Inhibitor Cocktail (1X), PMSF (10 mM)). Post-nuclear cell lysate was subjected to chloroform-methanol precipitation and the protein pellet was resuspended in 400 µL of blocking buffer (0.2 % S-methyl methanethiosulfonate (MMTS) (v/v); 5 mM EDTA; 100 mM HEPES; pH 7.4) and incubated for 15 min at 42°C. MMTS was removed by three rounds of chloroform-methanol precipitation and protein pellets were dissolved in 2SB buffer (2 % SDS; 5 mM EDTA; 100 mM HEPES; pH 7.4). 1/10 of each sample was retained as input control. For thioester bond cleavage and capture of free thiol groups, freshly prepared solution of neutral 2M hydroxylamine solution (pH 7.0-7.5, final concentration 400 mM) was added together with thiopropyl sepharose to each experimental sample. In negative control samples, same concentration of sodium chloride was used instead of hydroxylamine. After 1 h incubation, beads were collected, and the proteins were eluted by incubation in SDS sample buffer supplemented with 5mM DTT and analyzed by Western blotting.

### Immunoprecipitation

Cells were lysed in 1% DDM lysis buffer (1 % Dodecyl β-D-maltoside (DDM) in DPBS; 10 µM ML211; Phosphatase Inhibitor Cocktail 2 (1:100); Protease Inhibitor Cocktail (1X) and PMSF (10 mM)). 500 µg of lysate was used for each immunoprecipitation. 1/10 of each sample was retained as input control. For immunoprecipitation, lysates were incubated overnight at 4°C with 1 µg of antibody against the protein of interest. This was followed by incubation with Protein A agarose beads for 4 hours at 4°C. Immunoprecipitated proteins were eluted off beads by incubation with 1X Laemmli SDS sample buffer supplemented with 5 mM DTT for 15 min at 80°C with shaking. Eluted proteins were then analyzed by immunoblotting.

### Quantitative real-time PCR

Total RNA was prepared from T cells using TriZol reagent (Invitrogen). cDNA was synthesized using Superscript reverse transcriptase and oligo(dT) primers (Invitrogen) and gene expression was examined with a Bio-Rad iCycler Optical System using iQ™ SYBR green real-time PCR kit (Bio-Rad Laboratories, Inc.). The data were normalized to 18S. DHHC21: F-GGCTCTTTCTGCAGTTGTGT; R-GGCCGCGATATTTCTTCACA. T-bet: F-TGCGCCAGGAAGTTTCATTT; R-GGGCTGGTACTTGTGGAGAGACT. Gata3: F-GCAGCCTGCTGGGAGGAT; F-TAGAGGTTGCCCCGCAGTT. RORγt: F-AGCATCTATAGCACTGACGG; R-CAGAAACTGGGAATGCAGTG; RORα: F-TCTCCCTGCGCTCTCCGCAC; R-TCCACAGATCTTGCATGGA.

### In vitro differentiation and activation of naïve CD4^+^ T cells

Naïve T cell differentiation was performed as described previously (16). Naïve CD4+ T cells were isolated from murine spleens and lymph nodes using naïve CD4^+^ T cell purification kit (Miltenyi Biotech, Cat. 130-104-453) according to the manufacturer’s instruction. Isolated T cells were activated with anti-CD3 and anti-CD28 (BD Pharmingen, 2μg/ml) under Th0 (hIL-2 - Peprotech,50U/ml), Th1 (anti-IL-4 - Bioexcel,10μg/ml, IL-12 - Peprotech,10ng/ml, hIL-2–50U/ml), Th2 (anti-IFN-γ-Bioexcel,10μg/ml, IL-4-Peprotech, 10ng/ml, hIL-2–50U/ml), and Th17 (anti-IFN-γ-10μg/ml, anti-IL-4–10μg/ml, IL-6-Peprotech,20ng/ml, TGF-β-Peprotech,2ng/ml) polarizing conditions for 4 days. Then, cells were re-stimulated with anti-CD3/CD28 for 4 hrs. Supernatant was collected to assess cytokine production by ELISA (R&D Systems) and differentiated cells were used for mRNA expression analysis.

For T cell activation, WT or DHHC21^dep^ T cells were purified using CD4^+^ T cell isolation kit (Miltenyi Biotech, Cat. 130-104-453) and activated with plate-bound anti-CD3 and/or anti-CD28 (eBioscience). IL-2 production was determined 24 h after T-cell activation by ELISA according to manufacturer’s instructions (R&D Systems). CD69 production was analyzed by flow cytometry after 2 days of anti-CD3 and/or anti-CD28 stimulation.

### Flow cytometry

Splenocytes were harvested and single-cell suspensions were prepared in PBS/5% FCS buffer. Cells were kept on ice during all the procedures. For the extracellular markers, cells were stained with appropriate surface antigens. The following antibodies were obtained from BD eBioscience, or Invitrogen and were as follows: PerCP-Cy5.5-conjugated anti-CD3 (clone SK7, 1:20 dilution), CD4 PE (clone GK1.5, 1:200), CD8 Fitc (clone 5H10-1, 1:800), CD62L FITC (clone MEL-14, 1:800 dilution), CD44 APC (clone IM7, 1:500 dilution), CD69 PE (clone H1.2F3, 1:100 dilution). Live/dead assays were determined using the Aqua Dead Cell Stain Kit (ThermoFisher, Cat. L34957). Gating was determined using the Fluorescence minus one approach. Detection of cell surface markers was conducted using a Beckman-Coulter Gallios Flow Cytometer (BD Biosciences, San Jose, CA, United States) and data were analyzed by Kaluza Analysis Software, BD FACs Canto and FlowJo v. 10.6.2.

### Fura-2 Calcium Imaging

Cells were loaded with Fura-2 AM as described previously(30, 31) and placed in an imaging chamber with a glass bottom pre-coated with Poly-L-lysine. Images were taken on a Nikon TiS inverted microscope (Tokyo, Japan) with a 40× oil immersion objective, and images were taken every 3 seconds with a Photometrics Evolve electron-multiplying charged-coupled device camera (Tucson, AZ). Cells were pre-treated with αCD3 antibody and secondary IgG antibody was added after baseline recording to initiate TCR signaling.

## Acknowledgements

We thank Ethan Marin (Yale School of Medicine) for providing us with Zdhhc21^dep^ mouse strain.

This work was supported by startup funding from McGovern Medical School at University of Texas Health Science Center (to A.M.A.) and National Institute of General Medical Sciences grants R01GM115446 (to A.M.A.) and R01GM081685 (to D.B.).

